# Learned Visual Relevance Shapes Perceptual Inference: Evidence from Eyes to Hands

**DOI:** 10.64898/2026.08.30.748064

**Authors:** Kun Dong, Jan-Mathijs Schoffelen, Uta Noppeney

**Author notes:** Senior author (Uta Noppeney).

## Abstract

Perceptual inference requires the brain to decide which signals come from common causes and should hence be integrated or else be segregated. Leading accounts posit that sensory signals are weighted near-optimally by their relative precision consistent with Bayesian principles. It is unknown whether learnt behavioural relevance influences how the brain weighs signals in sensory integration.

This study investigated whether visual priority acquired via instrumental learning can reweight sensory inputs in a subsequent unrelated spatial localisation task. In the visual instrumental learning phase, participants responded to one of two differently coloured visual flashes presented in a cluttered environment while withholding their response to the other one. In the audiovisual phase, they located brief sounds presented synchronously midway between brief flashes of the previously relevant target and the competing non-target.

Observers’ perceived sound location shifted toward the previously relevant target as audiovisual disparity increased. Crucially, this audiovisual spatial bias arose dynamically across the entire cascade of perceptual inference and decision making: from rapid saccades and sustained gaze orienting towards the target location, both occurring after the target’s disappearance to initial deviations of manual response trajectories and on a fraction of trials, a change of mind with a return towards the sound as accumulating evidence suggested a separate cause structure.

Perceptual inference is thus governed not only by sensory precision but also by visual priorities acquired through instrumental learning. What we learn to prioritise in vision changes where we perceive sounds even in novel and unrelated contexts.

## Results

### Learned visual relevance biases auditory localisation towards the target

To form a coherent percept of the environment, the brain needs to infer the causes of the noisy sensory inputs. When visual and auditory signals occur in synchrony, the brain tends to integrate them weighted near-optimally by their relative precision into a unified percept consistent with Bayesian principles (Alais & Burr, 2004). This precision-weighted integration leads to audiovisual spatial biases (Meijer & Noppeney, 2020; Rohe & Noppeney, 2015). Because the visual signal is typically more precise than the auditory signal, observers perceive the sound shifted towards a synchronous yet spatially displaced flash – a perceptual illusion coined spatial ventriloquism. Traditionally, spatial ventriloquism has been regarded as a relatively automatic audiovisual integration process regardless of top-down endogenous spatial attention or bottom-up visual salience. Driver (1996) showed that displaced lip movements shifted the perceived location of speech. Bertelson et al. (2000) subsequently found that deliberate attention towards one visual event did not increase its influence over sound localisation. Similarly, Vroomen et al. (2001) showed that a visual singleton captured exogenous attention while the perceived sound was displaced towards competing visual elements elsewhere.

Recent research has shown that visual priority acquired via instrumental learning in one context influences subsequent visual selection and oculomotor dynamics in new unrelated contexts (Anderson et al., 2011; Hickey & van Zoest, 2012).

This study investigated whether prior instrumental learning can also influence how the brain weights and integrates sensory signals in perceptual inference and decision making.

Participants alternated between an instrumental visual detection phase and an audiovisual sound localisation phase (Figure 1A and 1B). In Phase A, they responded to a preassigned target-coloured blob (either orange or cyan) while withholding responses to a physically matched non-target. In Phase B, both blobs appeared simultaneously at equal distances on opposite sides of the sound. Both blobs remained task-irrelevant and differed primarily in their previously acquired behavioural relevance. Localisation errors (*Absolute bias*) were signed relative to the target side, such that positive values indicated a targetward shift. Across all tested audiovisual disparities, reported sound locations shifted consistently towards the targets (Figure 1C, left panel).

**Figure 1.**
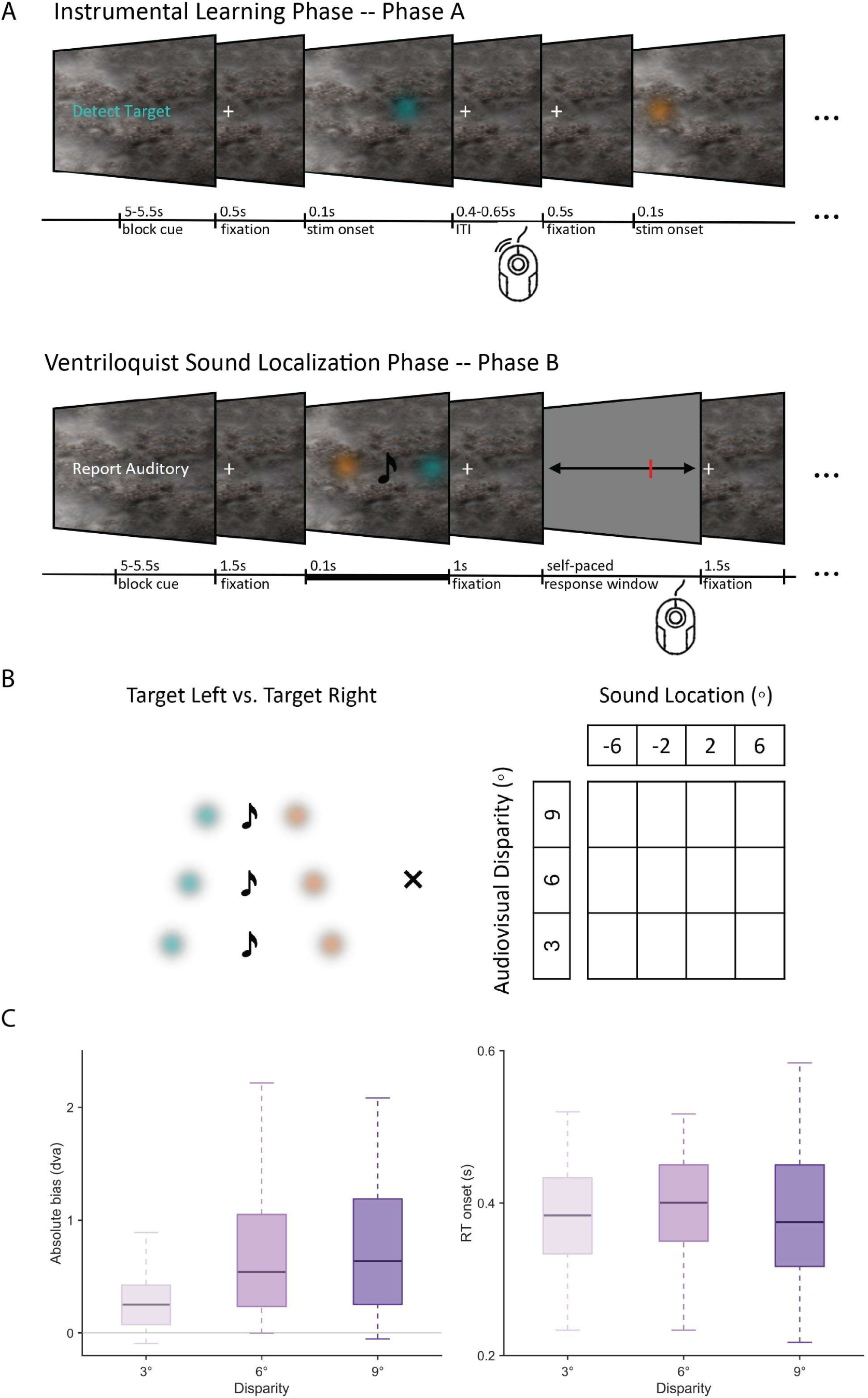
Experimental task, design and behavioural results. (A) Experimental task sequence. In Phase A, participants responded to the target while withholding responses to the non-target. In Phase B, participants localised brief sounds with the target and non-target competing. (B) Phase B stimulus geometry. Target location was counterbalanced, either left or right side of the sound. Sounds were presented at −6°, −2°, 2°, or 6° visual angle, with audiovisual disparities of 3°, 6°, or 9° visual angle. (C) Behavioural results in Phase B. Left: participant-level mean (*Absolute bias*) showing consistent targetward localisation bias in final reports. Signed so that positive values indicate localisation towards the learned target. Right: participant-level median *Response initiation latency*. Boxplots show participant-level summaries from the retained behavioural sample (n = 26); boxes indicate medians and interquartile ranges, and whiskers extend to 1.5× the interquartile range. Trial-level linear mixed-effects modelling showed that *Absolute bias* increased as audiovisual disparity increased (F(1, 25,653) = 5.35, *p* = 0.021), with no reliable quadratic effect (F(1, 25,653) = 2.21, *p* = 0.137). Model-estimated bias was greater than zero at 3° (0.35°, one-sided *p* = 0.0029), 6° (0.68°, *p* = 8.52 × 10−7), and 9° (0.81°, *p* = 3.19 × 10−8). Holm-corrected contrasts showed larger bias at 6° and 9° than at 3° (*p* = 4.58 × 10−5 and 1.17 × 10−8), whereas 6° and 9° did not differ reliably (*p* = 0.098). *Response initiation onset* also varied across audiovisual disparity, with reliable linear (F(1, 25,652) = 6.31, *p* = 0.012) and quadratic (F(1, 25,652) = 4.32, *p* = 0.038) effects.

A trial-level linear mixed-effects model showed that targetward localisation bias (*Absolute bias*) increased linearly as audiovisual disparity increased (F(1, 25,653) = 5.35, *p* = 0.021) without a reliable quadratic effect (F(1, 25,653) = 2.21, *p* = 0.137). Model-estimated bias was reliably greater than zero at 3° (0.35°, one-sided p = 0.0029), 6° (0.68°, one-sided p = 8.52 × 10^-7), and 9° (0.81°, one-sided *p* = 3.19 × 10^-8) visual angle. Holm-corrected contrasts confirmed the bias was larger at 6° and 9° than at 3° disparity, whereas the 6° and 9° conditions did not differ reliably. *Response initiation latency* also varied with audiovisual disparity (Figure 1C, right panel), showing reliable linear (F(1, 25,652) = 6.31, *p* = 0.012) and quadratic (F(1, 25,652) = 4.32, *p* = 0.038) effects.

### Targetward early oculomotor orienting during instrumental learning

Oculomotor dynamics during the instrumental task revealed that this visual priority emerged early. While small gaze biases do not necessarily explain attention allocation mechanistically, they provide a sensitive peripheral readout of attentional prioritisation (van Ede et al., 2019; Liu et al., 2022, 2025). In Phase A, both targets and non-targets evoked transient orienting and subsequent recentring (Figure 2). However, targets elicited a distinct early positive saccade rate bias towards the onset side (0.133 to 0.294 s poststimulus onset, cluster *p* = 0.001), followed by a later negative cluster (0.349 to 0.500 s, cluster *p* = 0.008). Non-targets also drove transient orienting, including positive clusters from 0.143 to 0.254 s (cluster *p* = 0.018) and from 0.548 to 0.658 s (cluster *p* = 0.048), as well as a negative cluster from 0.309 to 0.465 s (cluster *p* < 0.001). Crucially, the target-minus-non-target directional saccade rate contrast was positive from 0.131 to 0.328 s (cluster *p* < 0.001), revealing a strong early target-specific orienting bias.

**Figure 2.**
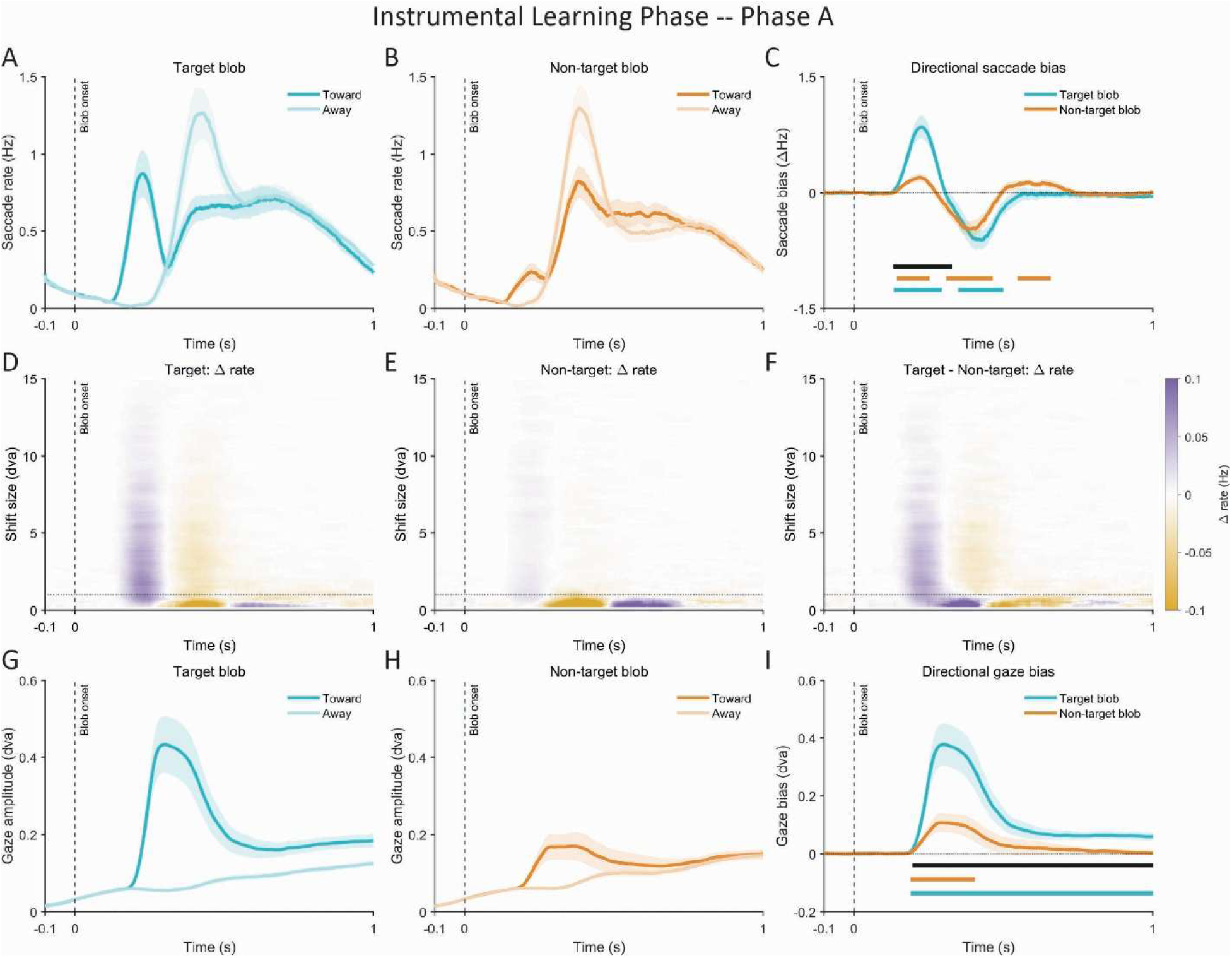
Early visual orienting during instrumental learning. (A) Phase A saccade rate results for targets, shown separately for shifts towards and away from the target. Traces show group means with SEM shading across participants (n = 25), and the dashed vertical line marks flash onset. (B) Phase A saccade rate results for non-targets, layout similar to (A). (C) Directional saccade rate bias, defined as towards minus away rate, for both targets and non-targets. Horizontal bars indicate significant cluster-corrected intervals from participant-level dependent-samples cluster-permutation tests over the 0–1 s poststimulus window; colour-matched bars refer to the corresponding conditions and the black bar refers to target-minus-non-target contrasts. Targets evoked an early positive bias from 0.133 to 0.294 s poststimulus onset (cluster *p* = 0.001), followed by a later negative cluster from 0.349 to 0.500 s (cluster *p* = 0.008). Non-targets showed positive clusters from 0.143 to 0.254 s (cluster *p* = 0.018) and 0.548 to 0.658 s (cluster *p* = 0.048), and a negative cluster from 0.309 to 0.465 s (cluster *p* < 0.001). The target-minus-non-target contrast showed a stronger early target-specific bias from 0.131 to 0.328 s (cluster *p* < 0.001). (D–F) Shift-size-by-time heatmaps showing signed saccade rate differences for targets (D), non-targets (E), and the target-minus-non-target contrast (F). Positive values indicate shifts towards the flash side. (G) Signed gaze displacement for targets, separated by towards and away directions. Traces show group means with SEM shading across participants (n = 25). (H) Signed gaze displacement for non-targets, separated by towards and away directions. (I) Directional gaze bias for targets and non-targets. Positive values indicate displacement towards the flash side. Gaze towards targets was positive from 0.190 to 1.000 s after blob onset (cluster *p* < 0.001), whereas non-targets showed a shorter positive cluster from 0.190 to 0.405 s (cluster *p* = 0.017). The target-minus-non-target signed-gaze contrast was positive from 0.196 to 1.000 s (cluster *p* < 0.001).

This bias persisted in continuous gaze displacement. Targets-elicited gaze displacement was positive from 0.190 to 1.000 s poststimulus onset (cluster *p* < 0.001), while non-targets produced a much shorter positive cluster (0.190 to 0.405 s, cluster *p* = 0.017). The target-minus-non-target signed-gaze contrast remained positive from 0.196 to 1.000 s (cluster *p* < 0.001).

### Targetward gaze bias persists during sound localisation in a cluttered multisensory environment

This target-driven oculomotor bias persisted in the multisensory environment of Phase B, where the target and non-target competed simultaneously around the brief sound (Figure 3). Saccade rate analyses revealed an early targetward bias from 0.173 to 0.299 s after audiovisual stimulus onset (cluster *p* = 0.016). This effect emerged despite closely matched overall rates of targetward and non-targetward saccades, indicating a brief but reliable early targetward bias during stimulus encoding. Continuous gaze displacement captured a more sustained targetward effect, with gaze displaced towards the target side from 0.229 to 1.000 s after audiovisual stimulus onset (cluster *p* < 0.001).

**Figure 3.**
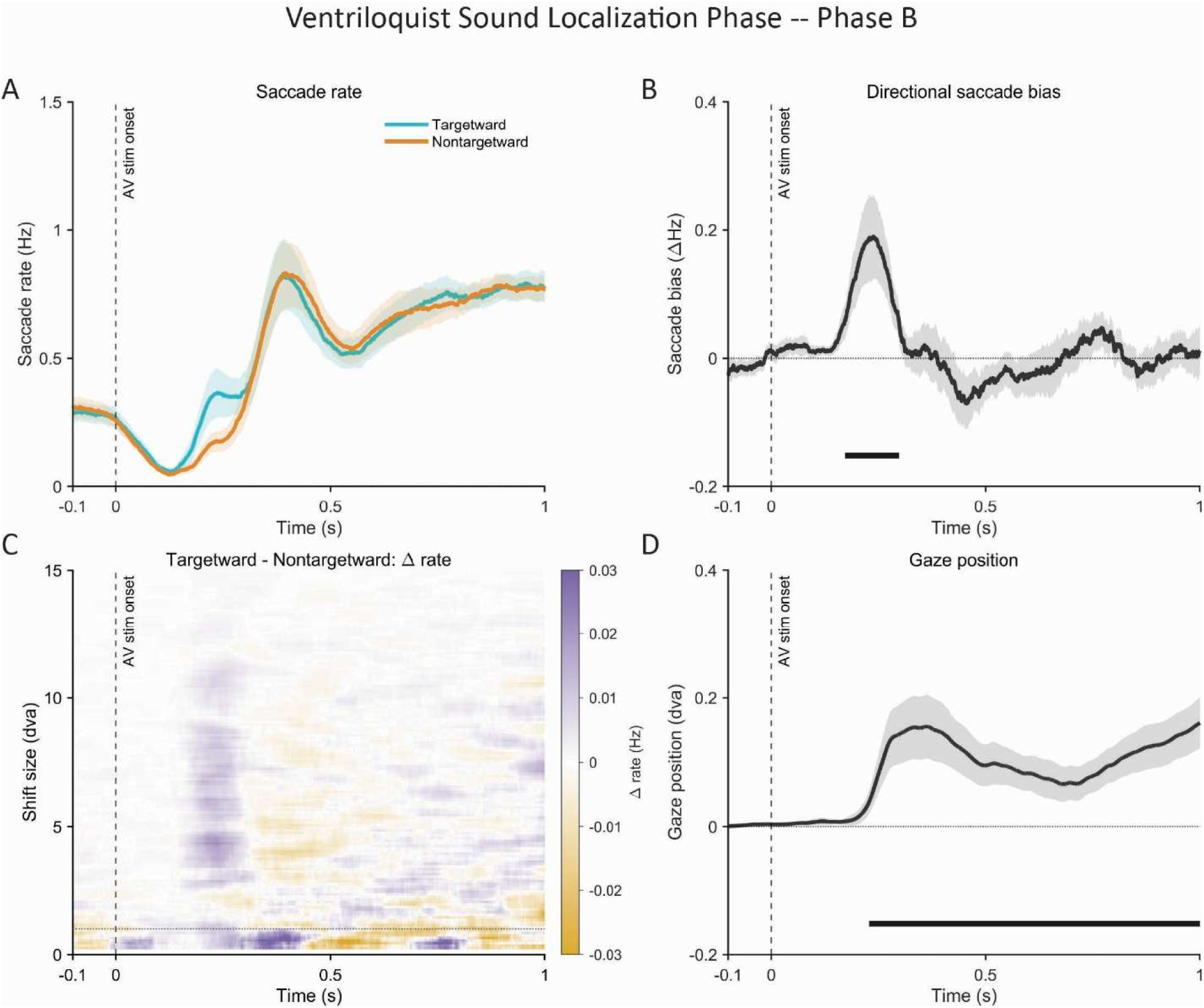
Targetward oculomotor dynamics during sound localisation. (A) Phase B saccade rates towards the target side and towards the non-target side post audiovisual stimulus onset. Traces show group means with SEM shading across participants (n = 25), and the dashed vertical line indicates audiovisual stimulus onset. (B) Directional saccade rate bias, defined as targetward minus non-targetward rate. Horizontal bars indicate significant cluster-corrected intervals from participant-level dependent-samples cluster-permutation tests over the 0–1 s post-stimulus window. The directional contrast revealed an early targetward cluster from 0.173 to 0.299 s after audiovisual stimulus onset (cluster mass = 356.28, cluster *p* = 0.016). (C) Shift-size-by-time heatmap for the targetward-minus-non-targetward saccade-rate difference. Positive values indicate a targetward pull. (D) Signed gaze position, so that positive values indicate gaze displacement towards the target side. Gaze was displaced towards the target from 0.229 to 1.000 s after audiovisual stimulus onset (cluster mass = 2443.93, cluster *p* < 0.001). Traces show group means with SEM shading across participants (n = 25), and a dashed vertical line indicates audiovisual stimulus onset.

### Motor trajectories and changes-of-mind reveal a targetward pull during decision formation

To trace whether this bias actively shaped decision formation before the final reports, we analysed continuous mouse trackball trajectories in Phase B. Motor trajectories provide a continuous readout of evolving decision dynamics, reflecting how the motor system actively constructs a choice as evidence unfolds (Konovalov & Krajbich, 2020; Grenke et al., 2025). Motor trajectories deviated towards the target across all audiovisual disparities (Figure 4A). Cluster-based permutation tests against zero confirmed positive targetward trajectory clusters at 3° from 19% to 100% of normalised movement progress, at 6° from 9% to 100%, and at 9° from 1% to 100% (all cluster *p* = 2.00 × 10−4).

**Figure 4.**
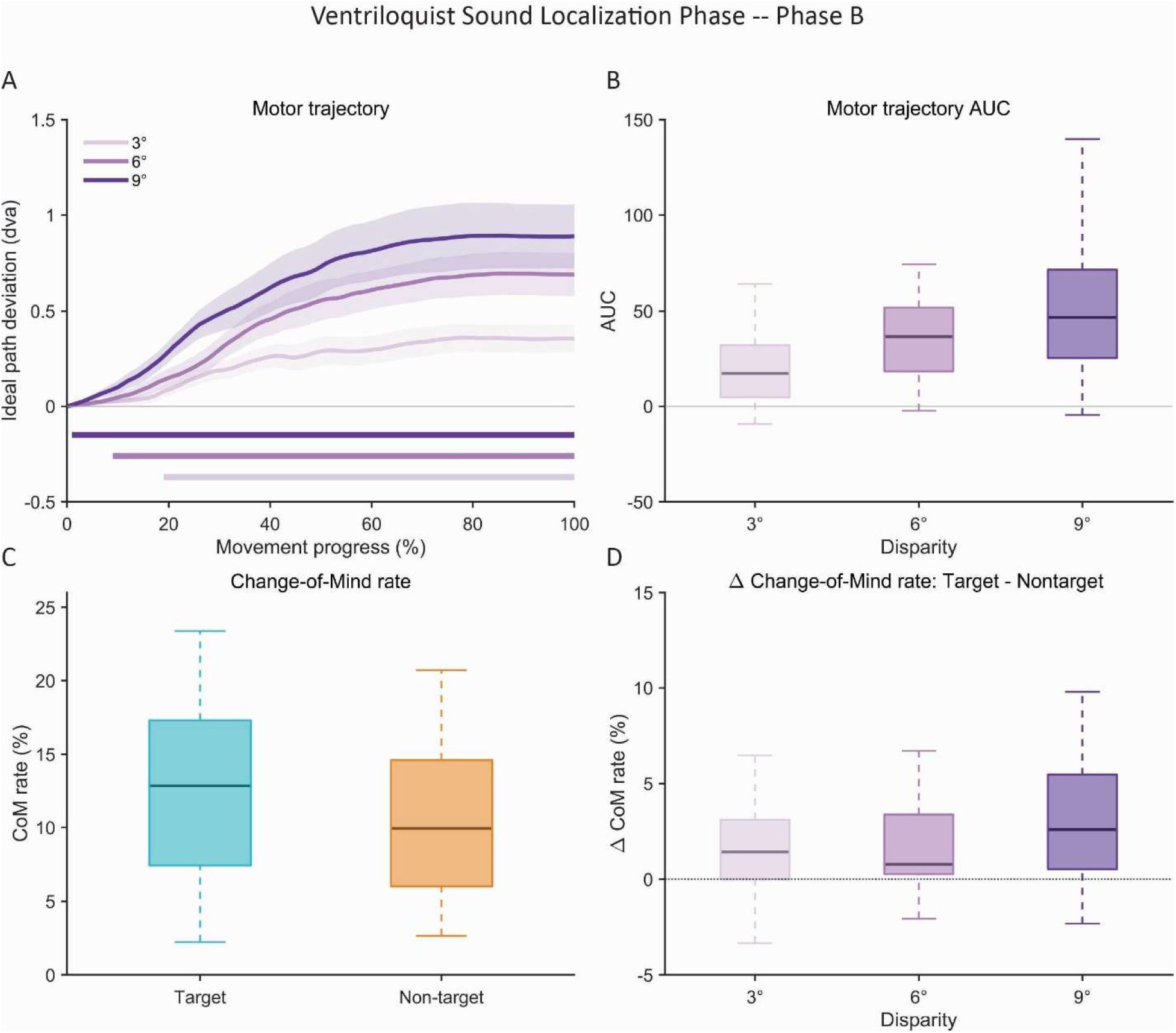
Motor trajectories and CoM reversals reveal a targetward pull in the motor system. (A) Group-averaged motor trajectory deviation across normalised movement progress across 3°, 6°, and 9° audiovisual disparities. Positive values indicate deviation towards the target side relative to a direct sound-reference movement path. Shaded ribbons show SEM across participants, and horizontal bars mark significant cluster-corrected intervals in which trajectories differed from zero. Trajectories were targetward at all disparities, with positive clusters from 19–100% of movement progress at 3°, 9–100% at 6°, and 1– 100% at 9° (all cluster *p* = 2.00 × 10−4). (B) Participant-level motor trajectory area under the curve (*AUC*) by disparity. Positive values indicate targetward trajectory area. Boxplots show participant-level summaries from the retained motor sample (n = 26); boxes indicate medians and interquartile ranges, and whiskers extend to 1.5× the interquartile range. The selected mixed-effects model showed a linear effect of audiovisual disparity (F(1, 50) = 35.73, *p* = 2.37 × 10−7) and no reliable quadratic effect (F(1, 50) = 0.20, *p* = 0.661). Model-estimated *AUC* was greater than zero at 3° (23.61, one-sided Bonferroni-corrected *p* = 0.014), 6° (44.98, one-sided Bonferroni-corrected *p* = 1.14 × 10−5), and 9° (61.50, one-sided Bonferroni-corrected *p* = 3.17 × 10−8). Bonferroni-corrected pairwise contrasts further showed an increase in targetward area across disparities (all corrected p ≤ 0.036). (C) Participant-level *CoM* rates for target-directed and non-target-directed revisions. Target-directed reversal events were more frequent than non-target-directed reversal events (12.70% ± 1.25% SEM versus 10.32% ± 1.00% SEM; t(25) = 4.21, one-sided *p* = 1.42 × 10−4). This targetward advantage was reliable within each disparity after one-sided Bonferroni correction: 3° (*p* = 0.0086), 6° (*p* = 0.0054), and 9° (*p* = 3.76 × 10−4). (D) *ÄCoM* rate by disparity, defined as target-directed minus non-target-directed *CoM* rate. Positive values indicate more target-directed than non-target-directed reversals. The winning mixed-effects model showed a linear effect of audiovisual disparity (F(1, 50) = 4.86, *p* = 0.032) and no reliable quadratic effect (F(1, 50) = 1.37, *p* = 0.248). Model-estimated *ΔCoM* rate was greater than zero at 3°, 6°, and 9° after one-sided Bonferroni correction, whereas pairwise disparity differences were not reliable after correction for all pairs. Boxplots show participant-level summaries from the retained motor sample (n = 26). See also Figure S2.

We summarised this deviation using the area under the curve (*AUC*; Figure 4B), a standard mouse-tracking measure that captures the magnitude and direction of deviation from a reference path (Pfister et al., 2024). The winning linear mixed-effects model showed a strong linear effect of audiovisual disparity on the *AUC* (F(1, 50) = 35.73, *p* = 2.37 × 10−7), with no reliable quadratic effect (F(1, 50) = 0.20, *p* = 0.661). Model-estimated *AUC* was reliably greater than zero at 3° (23.61, one-sided Bonferroni-corrected *p* = 0.014), 6° (44.98, one-sided Bonferroni-corrected *p* = 1.14 × 10−5), and 9° (61.50, one-sided Bonferroni-corrected *p* = 3.17 × 10−8). Bonferroni-corrected pairwise contrasts further showed increasing targetward *AUC* across audiovisual disparities (all corrected *p* ≤ 0.036).

Change-of-mind (CoM) reversals, which capture within-trial revision of an evolving decision (Resulaj et al., 2009; van den Berg et al., 2016; Vivar-Lazo & Fetsch, 2026), provide a converging readout (Figure 4C and 4D). Target-directed reversal events were more frequent than non-target-directed events (12.70% ± 1.25% SEM versus 10.32% ± 1.00% SEM, t(25) = 4.21, one-sided *p* = 1.42 × 10−4). This target-dominant reversal was reliable within each audiovisual disparity after one-sided Bonferroni correction: 3° (*p* = 0.0086), 6° (*p* = 0.0054), and 9° (*p* = 3.76 × 10−4).

For *ΔCoM*, mathematically defined as target-directed minus non-target-directed CoM rate, the winning linear mixed-effects model showed a linear effect of audiovisual disparity (F(1, 50) = 4.86, *p* = 0.032), with no reliable quadratic effect (F(1, 50) = 1.37, *p* = 0.248). Model-estimated *ΔCoM* was greater than zero at 3°, 6°, and 9° audiovisual disparities after one-sided Bonferroni correction, though pairwise disparity contrasts were not statistically reliable. Ultimately, participants were more often initially biased towards the target before redirecting their response towards the perceived sound, demonstrating that learned visual relevance actively shaped the unfolding motor decision before final commitment.

## Discussion

By combining psychophysics, eye-tracking, and continuous motor trajectories, we show that instrumentally learned visual priority biased multisensory perceptual inference from early orienting to final manual responses. When a learned target and a physically matched non-target competed for binding with the same sound, the perceived sound location shifted towards the learnt target, progressively for increasing audiovisual disparity. The same targetward bias appeared dynamically across the entire cascade of perceptual inference and decision making: from rapid saccades and sustained gaze orienting towards the target location, both occurring after the target’s disappearance, to initial deviations of manual response trajectories and on a fraction of trials, a change of mind with a return towards the sound as accumulating evidence suggested a separate cause structure. Collectively, our results show that learned visual priority does not merely bias a late reporting process, but alters the weighting of sensory inputs during perceptual inference.

Our findings challenge the classic Bayesian and maximum likelihood accounts of multisensory integration, where signals are weighted purely by their relative precision (Ernst & Banks, 2002; Alais & Burr, 2004; Rohe & Noppeney, 2015; Meijer & Noppeney, 2020). Two physically matched visual signals at equal spatial disparity exerted different influences on observers’ perceived sound location depending on the relevance they had acquired through prior instrumental learning in an unrelated context. This profile is unlikely to be explained via attentional mechanisms because previous research showed that neither directing attention to one visual event nor capturing attention with an exogenous visual singleton influenced sensory weighting in multisensory integration (Bertelson et al., 2000; Vroomen et al., 2001). They move beyond previous evidence that learned selection history can shape visual priority and oculomotor behaviour (Anderson et al., 2011; Hickey & van Zoest, 2012) by showing that such learned priority also reweights input in multisensory perception and decision making. Future studies are needed to investigate whether Pavlovian conditioning with different reward values can similarly influence the weighting of sensory inputs in perception (Cheng et al., 2020; Vakhrushev & Pooresmaeili, 2024).

Oculomotor dynamics reveal the emergence of visual priority. During the instrumental learning phase, target flashes elicited a stronger early saccade bias than non-target flashes, followed by a sustained targetward gaze displacement. This targetward bias persisted into the subsequent sound localisation task when the same target flashes were no longer task-relevant. Targets again elicited rapid saccades and drove gaze displacement, thereby serving as a time-resolved readout of active attentional selection (van Ede et al., 2019; Liu et al., 2022, 2025). Critically, however, because the flashes lasted only 100 ms, the saccades and gaze biases did not alter the visual inputs, and yet they were predictive of the later sound biases.

Continuous manual trajectories extended this targetward pull from sensory orienting during instrumental learning and sensory encoding to decision formation. Movements deviated toward the target across all audiovisual disparities, and participants more frequently initiated movement toward the target before redirecting to the perceived sound location than with non-targets. Continuous trajectories reveal evolving decision states that remain hidden before the final response (Konovalov & Krajbich, 2020), while within-trial reversals capture decision revisions after movement initiation (Resulaj et al., 2009; van den Berg et al., 2016). Here, the target did not merely bias sound localisation in final reports. It actively shaped the movement trajectory that provided insights into the evidence accumulation process.

This pattern supports an embodied view of perceptual inference (Lepora & Pezzulo, 2015; Verdonck et al., 2021; Balsdon et al., 2023). From a strict serial processing perspective, sensory evidence accumulates until a decision bound is reached, and the completed choice is then passed to the motor system for execution. By contrast, the combined evidence of early oculomotor orienting, motor trajectory deviation, and target-driven reversals (i.e. change of mind) demonstrate that evolving motor actions carry information about the dynamically evolving perceptual decision (Gherman & Philiastides, 2015; van den Berg et al., 2016; Balsdon & Philiastides, 2024). Recent neurophysiological work in non-human primates further supports this by demonstrating that choice and confidence signals can be updated concurrently within sensorimotor neural populations (Vivar-Lazo & Fetsch, 2026). Collectively, these findings indicate that the motor system is not simply the endpoint of a completed perceptual choice. The unfolding motor dynamics provided a continuous readout of how learned visual relevance influences perceptual inference and decision making before final commitment.

More broadly, adaptive behaviour requires the brain to infer not only which sensory signals belong together, but also how to weigh them optimally into a coherent percept of the environment. Our results show that visual priority persisted beyond the instrumental task and influenced sound localisation, oculomotor dynamics, and manual trajectories. Learned visual relevance therefore provides a route through which previous experience shapes how the brain dynamically weights sensory inputs for perceptual inference, decision making and report in a cascade of processing from early saccades to initial deviations of manual response trajectories and final read out.

## Methods

### EXPERIMENTAL MODEL AND STUDY PARTICIPANT DETAILS

#### Experimental design and sequence

The experiment investigated how instrumentally learned relevance assigned to a coloured (either orange or cyan, counterbalanced across participants) visual flash affects subsequent sound localisation in a cluttered multisensory environment. Participants completed a two-day experiment alternating between an instrumental visual detection task (Phase A) and a subsequent audiovisual sound localisation task (Phase B). Each day began with a long Phase A block, followed by an alternating sequence yielding 65 phases per day, leading to 130 phases across the whole experiment.

Phase A established and maintained the behavioural relevance of the assigned coloured visual flash (i.e., the target). Participants were instructed to detect the target as quickly and accurately as possible, while withholding responses to the non-target. Phase A established a learned priority for the target through instrumental learning and its relevance to subsequent multisensory perceptual decision making (see Figure S1 for an overview of Phase A behavioural performance).

Phase B tested how this learned priority influences subsequent sound localisation in a cluttered audiovisual environment. On each trial, participants localised a brief sound with two competing visual flashes (i.e., target vs. non-target) presented in synchrony. The two visual flashes surrounded the sound source at equal distances, creating a competing integration dynamic over observers’ audiovisual integration processes. Participants localised the sound on a continuous horizontally displaced response bar using a mouse trackball (see also Figure 1 for visual demonstration).

Combined with this specific setup and concurrent eye-tracking, it provided a time-resolved measure of oculomotor orienting towards the visual flashes. The mouse trackball provided a variety of readouts during the action formation procedure, including final sound reports, response initiation latency, motor trajectory, and change-of-mind reversals. Together, these measures allowed us to ask not only whether the visual priority biases audiovisual integration, but also how this bias became visible across oculomotor dynamics, action formation, and final reports.

#### Participants

Twenty-seven adult healthy human volunteers participated in the study (19 female, 8 male; mean age 25.6 years, SD 4.8 years, range 18-38 years). One participant dropped out midway during the experiment, leaving twenty-six participants for analysis. Of the participants included in this study, twenty-five were right-handed, and one was left-handed, according to the Edinburgh Handedness Inventory (Oldfield, 1971). All participants reported normal or corrected-to-normal vision, normal hearing, and no history of neurological or psychiatric conditions. All volunteers provided written informed consent and were naïve to the study’s purposes. They received monetary reimbursement for their participation in the experiment. The study was approved by the ethical board of CMO Arnhem/Nijmegen under registration number CMO2014/288 and was conducted in accordance with the principles outlined in the Declaration of Helsinki. Similar sample sizes have been reported across studies to successfully capture the dynamics of oculomotor orienting (van Ede et al., 2019; Liu et al., 2022).

#### Experimental setup

The presentation scripts were written with Psychtoolbox 3.0.19 running under MATLAB R2023a on a Windows computer. We used a Dell U3014 LED monitor with a 60 Hz refresh rate, a 64 × 40 cm screen size, and a resolution of 2560 × 1600 pixels for experimental presentation. Participants were comfortably seated in a dimly lit behavioural booth at a viewing distance of 60 cm. Auditory stimuli were delivered through Sennheiser HD 280 PRO headphones at 48,000 Hz. Visual rendering, including the dynamic background used through the main tasks, was controlled through OpenGL functions via Psychtoolbox. Button presses in Phase A for detection and continuous localisation motor trajectory were produced using a Kensington Orbit wireless trackball (K70992WW) using Psychtoolbox mouse polling. Eye-tracking calibration and event-message communication were controlled via the Eyelink Toolbox (Cornelissen et al., 2002).

#### Preliminary unisensory familiarisation and calibration tasks

Before the main experiment, participants completed unisensory familiarisation and calibration tasks. These tasks familiarised participants with the spatial mapping for both sensory modalities (i.e., visual and auditory), the response bar, and the mouse trackball. They also provided baseline estimates of sound localisation and participant-specific response-related variability (i.e., motor noise), independent of the main tasks.

Participants first completed a sound familiarisation task. Spatialised sounds were presented from different locations along the azimuth, with a green visual marker indicating the true location on the response bar. Participants did not need to respond during this task.

Participants then completed a unisensory sound task. On each trial, a brief spatialised sound was presented for 100 ms, and participants reported the perceived location on a response bar mapped to the stimulus range (from −15° to 15° visual angle). Participants responded using a mouse trackball at their own pace, with no fixed response deadline. To minimise motor planning/preparation biases, we randomised the initial cursor position on each trial. Following the final reports, the real sound location was displaced on the response bar as visual feedback. We later calculated participant-specific sound localisation estimates from this task as baselines.

Finally, participants completed a visual-motor calibration task using the same response setup. On each trial, we briefly presented a red vertical bar for 100 ms at a location sampled along the azimuth within the same range as mentioned above. After a 1-s interval, participants recalled the visual location using the response bar. No feedback was provided. This task was included to estimate participant-specific response noise, referred to as motor noise in this paper, which captures variance introduced by the response system itself, including motor variability and short-term memory demands (for a similar measurement of motor noise, see Hong et al., 2022).

#### Main experimental sequence

The main experiment comprised two experimental days with two alternating phases: an instrumental visual detection task (Phase A) and a subsequent audiovisual sound localisation task (Phase B). On each day, participants completed one long Phase A block, following an experimental sequence of A – (B – A) × 32, for 65 phases in total. The full experiment comprised 130 phases across the two testing days.

#### Phase A: Instrumental visual detection task

We used an instrumentally speeded visual detection task to establish and maintain the target’s priority. We instructed participants to respond to the targets as quickly and accurately as possible while withholding responses to non-targets. Both targets and non-targets were coloured visual flashes (orange or cyan) preassigned to participants, counterbalanced across participants, and remained fixed throughout the experiment. Those flashes were Gaussian blobs presented for 100 ms against a dynamic Shepard-zooming background, with a size of 2° visual angle. To equate perceptual lightness across colours, we transformed the seed for both colours into CIE L*a*b* space (Melgosa et al., 1994), matched it to the same target L* value, then converted it back to sRGB while preserving hue as far as the display gamut allowed.

The dynamic Shepard zooming background was generated using multilayer spectral texturing and rendered in real time using OpenGL functions via Psychtoolbox in MATLAB on a standard Windows PC with an NVIDIA Quadro K600 GPU, enabling dedicated smooth rendering during the experiment presentation (for a detailed description of the Shepard zooming background, see Berger, 2003; for a demonstration, see Psychtoolbox/PsychDemos/OpenGL4MatlabDemos/ShepardZoomDemo). Spectral texturing models the statistical relationships between bands of the texture’s spatial spectrum, resulting in realistic textures with an overall amplitude spectrum close to 1/f, mimicking the amplitude spectrum of natural images (Simoncelli & Olshausen, 2001). Naïve observers reported the stimuli as heavy clouds moving towards or away from them; for a similar setup in a previous study in our lab, see Conrad et al., 2013. This dynamic background created a semi-naturalistic cluttered visual environment throughout the whole experiment (across both phases) against which the visual flashes were presented.

There were a total of 31 possible locations along the azimuth of the visual flashes within the stimulus range (from −15° to 15° visual angle). At the beginning of each block, a written instruction of “Detect Target” was presented in the colour of the target for 5-5.5 s. Each trial began with a 0.5 s fixation period, during which a white fixation cross was presented at the centre of the screen. A visual flash was then presented briefly for 100 ms, with the fixation cross temporarily removed. Participants responded only when detecting a target. The inter-trial interval (ITI) varied pseudo-randomly between 400 and 650 ms, following a Poisson distribution with λ = 18 and a scaling factor of 20 to reduce temporal predictability (Lewis & Noppeney, 2010). At the end of each block, participants received performance-based feedback. The long Phase A block contained 124 trials, followed by 32 short blocks, each containing 31 trials. Participants hence each completed 2,232 trials across the whole experiment.

#### Phase B: Audiovisual sound localisation task

Following each Phase A block, participants completed an audiovisual sound localisation task. We used this task to test whether the object priority established via instrumental learning persisted and biased sound localisation in a cluttered multisensory setup (the same Shepard zooming background was retained in this phase).

Auditory stimuli were spatialised pink noises with a duration of 100 ms and 5 ms onset and offset ramps. We generated the spatialised materials by convolving the pink noises with spatially selected head-related transfer functions derived from the KEMAR dummy-head database at the MIT Media Laboratory (Gardner & Martin, 1995). Head-Related Transfer Functions (HRTF) from the database locations were interpolated to obtain the desired locations. Phase B included four sound locations along the azimuth with no elevation manipulation: −6°, −2°, 2°, and 6° visual angle. Visual flashes were presented surrounding the sound location in synchrony. In regular trials, visual flashes were presented at equal distances from the sound, with 3 levels of audiovisual disparity at 3°, 6°, or 9° visual angle. Target sides relative to the sound were balanced (either left or right). Therefore, Phase B followed a 4 × 3 × 2 factorial design, with four auditory locations, three audiovisual disparities, and two target sides.

At the beginning of each Phase B block, participants localised the sound source as accurately as possible. Each trial began with a 1.5 s fixation period. The sound and two visual flashes were then presented in synchrony for 100 ms, with the fixation cross temporarily removed. After audiovisual stimulus offset, a 1 s poststimulus fixation period followed. Participants then reported the perceived sound location using the mouse trackball on a response bar. Responses were self-paced, with no fixed response deadline, and no feedback was provided. Each block consisted of 20 trials: 18 regular trials and 2 catch trials. Catch trials were generated by pseudo-randomly changing the sound location of randomly selected regular trials to a different location while leaving the configuration of visual flashes unchanged. These trials were included to promote sustained spatial attention and reduce the likelihood that participants would solve the task by relying on underlying stimulus configuration geometries. We did not include catch trials in the formal analysis. Participants completed 32 Phase B blocks per day, for a total of 1,280 Phase B trials across the experiment.

#### Eye-tracking acquisition and gaze preprocessing

We consistently recorded the left eyes across all participants using an Eyelink 1000 Plus Eye Tracker (SR Research). Recordings started after applying a five-point calibration routine (HV5), followed by drift correction at the beginning of each block. Full recalibration was performed when necessary. Most recordings were made at the sampling rate of 1,000 Hz, whereas a small number of sessions were made at 2,000 Hz. Gaze data were preprocessed offline using the PuPl toolbox (Kinley & Levy, 2022), with dependencies including the Edf2Mat toolbox (https://github.com/uzh/edf-converter). Invalid pupil samples were trimmed before blink detection, and blinks were identified using PuPl’s built-in noise-based blink-detection procedure (Hershman et al., 2018). Samples surrounding each blink were removed with a 100-ms padding window before and after the blink. We did not perform gaze interpolation or apply additional smoothing. To place all recordings on a common temporal scale, we retained 1,000-Hz recordings at their native sampling rate, and downsampled 2,000-Hz recordings to 1,000 Hz. Epochs for subsequent oculomotor analyses were then extracted from the preprocessed gaze data using the task design files and Eyelink event messages. Recordings were visually inspected after preprocessing. One participant was excluded from the subsequent oculomotor analyses because more than 50% of eye-tracking samples were missing after preprocessing, resulting in a total of twenty-five participants included in the oculomotor analysis.

#### Response acquisition and trajectory recording

Detection responses in Phase A and localisation responses in Phase B were acquired using a Kensington Orbit wireless trackball. In Phase A, participants responded to targets by pressing the left trackball button. Responses were logged relative to the trial sequence and used to classify target detections, misses, and false alarms. In Phase B, the same device was used for final reports and to record the evolving motor trajectory leading to that final click. After the 1 s poststimulus fixation interval, a response bar was presented. On each trial, to minimise motor preparation/planning, the response-guidance cursor was placed at a random position on the bar. Participants navigated the cursor along the response bar using the trackball and clicked the left button to commit to the perceived sound location. Responses were self-paced, with no hard deadline for responses or feedback. We detected response initiation latency online from the continuous cursor stream. Movement onset of the trackball was defined as the first time point at which three consecutive response speed estimates exceeded 300 pixels/s. This measure was used to assess commitment readiness, separately from motor trajectory and final reports.

### QUANTIFICATION AND STATISTICAL ANALYSIS

We analysed all behavioural, oculomotor, and motor trajectory data using custom MATLAB and R pipelines. Gaze preprocessing was performed using the PuPl toolbox (Kinley & Levy, 2022); time-resolved cluster-based permutation tests were conducted with FieldTrip (Oostenveld et al., 2011); and linear mixed-effects modelling was performed in RStudio using R 4.4.3. The main R packages used for mixed-effects modelling and post hoc contrasts were lme4 (Bates et al., 2015), lmerTest (Kuznetsova et al., 2017), and emmeans (Lenth, 2023). We analysed continuous dependent variables using linear mixed-effects models, with participants as a random intercept. We treated audiovisual disparity as the main within-participant variable for Phase B behavioural and motor trajectory analyses. When relevant, candidate models included additional quadratic disparity terms (i.e., the square of audiovisual disparity), initial cursor position, and participant-specific motor noise estimated from the visual-motor calibration task. We fitted candidate models using maximum likelihood and compared them using the Akaike information criterion (AIC). We then refitted the winning model using restricted maximum likelihood for inference. Estimated marginal means were used for post hoc contrasts at 3°, 6°, and 9° audiovisual disparity. Pairwise contrasts and tests against zero as the baseline were corrected for multiple comparisons. Time-resolved gaze-position, saccade rate, and motor trajectory analyses were performed at the participant level using cluster-based permutation tests.

#### Behavioural sound localisation analysis

We examined whether target priority persisted in the audiovisual sound localisation phase using two behavioural metrics: *Absolute bias* and *Response initiation latency*.

To account for individual differences in baseline sound localisation, we first estimated participant-specific reference locations from the unisensory sound localisation task. For each participant, we regressed the reported sound location on the real location. The predicted sound reference for each participant

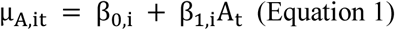

Where μ_A,it_ is the sound reference for participant i on trial t, A_t_ is the physical sound source of the given trial, and β_0,i_ and β_1,i_ are the participant-specific intercept and slope from the unisensory sound-localisation linear fit.

*Absolute bias* was mathematically defined by signing the localisation error according to the side of the target in the audiovisual sound localisation task:

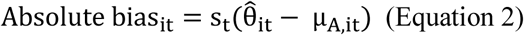

Where θ^_it_ is the final report of the perceived sound location and s_t_ is the target-side sign of the given trial, with s_t_ = 1 when the target was to the right of the sound and s_t_ = − 1 when the target was to the left. Positive values therefore indicate targetward localisation. We analysed *Absolute bias* and *Response initiation latency* as functions of audiovisual disparity and its quadratic effects. We also included the quadratic term to capture potential non-linearity in localisation biases across spatial disparity (i.e., an inverted-U shape; see also Cao et al., 2019). We defined *Response initiation latency* as the first time point after response window offset at which three consecutive response-speed estimates of the trackball exceeded 300 pixels/s. We excluded trials with *Response initiation latency* below 100 ms because of premature responses.

#### Oculomotor analysis

Oculomotor analyses tested whether target priority was visible from instrumental learning (Phase A) to sensory encoding (Phase B). Since all task-relevant manipulations were along the azimuth, we focused on horizontal oculomotor dynamics for subsequent saccade and gaze analyses. Gaze position was converted from pixels to degrees of visual angle using the display geometry.

In Phase A, gaze epochs were time-locked to flash onsets and baseline corrected using the 0-200 ms prestimulus interval. Gaze position was signed according to the side of the flash, with positive values indicating gaze displacement toward the flash and negative values indicating displacement away from the flash. We examined whether both target and non-target trials evoked reliable gaze orienting, and whether orienting differed between target and non-target trials.

In Phase B, gaze epochs were time-locked to audiovisual stimulus onset and baseline corrected using the 0-500 ms prestimulus interval. On each trial, gaze position was aligned with the sound location and then signed to the side of the target, with positive values indicating gaze displacement toward the target and negative values indicating displacement toward the nontarget. We examined whether gaze displacement was targetward during sensory encoding in the audiovisual sound localisation task.

In addition to gaze analysis, we quantified saccade dynamics. Saccades were detected using a velocity-based procedure adapted from previous oculomotor analyses of spatial attention studies (van Ede et al., 2019; van Harmelen et al., 2026). On each trial, gaze velocity was estimated as the temporal derivative of gaze position. The absolute velocity trace was smoothed with a 7 ms Gaussian-weighted moving window, and candidate events were identified when the smoothed velocity exceeded five times the trial-wise median absolute velocity. To avoid duplicate counting, a 100 ms refractory period was imposed after each detected event.

For each detected saccade event, gaze-shift magnitude and direction were estimated by comparing mean gaze position in the 50 ms interval before threshold crossing with mean gaze position in the 50–100 ms interval after threshold crossing. Events with an estimated magnitude below 0.075° visual angle were excluded as the movement was smaller than 1% of the unilateral stimulus range (i.e., 7.5° visual angle). We deliberately did not apply an upper amplitude cut-off to reflect the nature of orienting towards flashes. In Phase A, saccades were coded as either towards or away from the flash. In Phase B, saccade shifts were coded as targetward or non-targetward. Saccade rate time courses were computed using a 100 ms sliding integration window. Time-resolved analyses of gaze position and saccade rate were performed over the 0– 1 s post-stimulus onset time window. Statistical inference was based on participant-level time courses using cluster-based permutation tests.

#### Motor trajectory analysis

Motor trajectories were analysed to quantify how perceptual decision-making evolved before final commitment. Cursor tracking provides a time-continuous measure of decision dynamics and reveals information about latent decision states that is not always evident in final responses alone (Konovalov & Krajbich, 2020; Grenke et al., 2025). We focused on the horizontal cursor position, similar to oculomotor analysis. We only analysed regular trials in Phase B. On each trial, cursor position was converted from pixels to degrees. The motor trajectory was then expressed over 101 equally spaced movement-progress samples, corresponding to 0-100% of the decision making procedure. This allowed trajectories with different durations to be compared on a common movement-progress axis, following standard time-normalisation procedures in mouse-tracking analyses (Pfister et al., 2024). Linear interpolation was used for trajectory area analyses.

To depict unfolding targetward bias, we compared each trajectory with an ideal straight reference path from the initial cursor position to the participant-specific sound reference location, estimated from the unisensory sound localisation task (i.e., the participant’s perceived sound location) as mentioned above. Target-aligned trajectory deviation was then mathematically defined as:

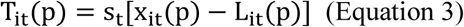

Where T_it_(p) is the target-aligned deviation for participant i, trial t, and movement progress sample p; x_it_(p) is the cursor position of the given trial at sample p; L_it_(p) is the ideal path from the initial cursor position to the sound reference at sample p; and s_t_ is the target-side sign, with s_t_ = 1 when the target was on the right side of the sound and s_t_ = − 1 when the target was on the left side. Positive values therefore indicate that the unfolding procedure moved towards the target, while negative values indicate the trajectory moved towards the non-target.

The primary scalar metric of the motor trajectory was the targetward area under the curve (*AUC*). *AUC* is a standard mouse-trajectory measure that summarises the magnitude and direction from an ideal or reference path (Pfister et al., 2024). Positive *AUC* values indicate that the response trajectory was pulled towards the target side in sum, whereas negative values indicate deviation away from the target (i.e., non-targetward). We analysed *AUC* using linear mixed-effects models, as described in the Behavioural sound localisation analysis section. Candidate models included linear and quadratic effects of audiovisual disparity; additional models included initial cursor position and participant-specific motor noise, estimated from the visual-motor calibration task. Model fitting and post hoc contrasts followed the general statistical approach described above.

Time-resolved trajectory effects were analysed using the full target-aligned movement deviation progress course (T_it_(p)). Participant-level trajectory time courses were examined with cluster-based permutation tests to identify periods during which trajectories were reliably biased towards or away from the target.

#### Change-of-mind analysis

Changes-of-mind (CoMs) were analysed to test whether the localisation trajectory shifted towards flashes before reversing back to the perceived sound location. CoMs have been used to characterise within-trial revisions of an evolving decision after an initial choice has already been expressed (Resulaj et al., 2009; van den Berg et al., 2016; Vivar-Lazo & Fetsch, 2026). For this analysis, we transformed motor trajectories into a sound-centred, target-coded coordinate frame. Similar to other analyses in this paper, positive movement indicated motion toward the target, and negative movement indicated motion away from the target. Motor trajectories were normalised to 101 movement-progress samples (i.e., 0-100%) using shape-preserving interpolation and smoothed with a Gaussian kernel with reflected edge padding. We ignored the first and last 5% of the movement progress to avoid classifying initial motor jitter or terminal correction as a change-of-mind. Candidate initial and reversal phases had to last at least 10% of the movement progress.

We scaled detection thresholds by participant-specific motor noise. The initial-excursion and reversal thresholds were set to 75% of motor noise and constrained to 0.30–3.00° visual angle. The return threshold was set to 50% of motor noise and constrained to 0.15–1.50° visual angle. A target-directed CoM event was identified when the cursor first showed a reliable excursion towards the target and then reversed towards the sound reference. Non-target-directed CoM events were detected using the mirrored trajectory (see also Figure S2 for visual illustration of CoM detection logic). For each participant and audiovisual disparity, we computed target-directed and non-target-directed reversal rates. The primary CoM measure was their difference, i.e., target-directed CoM minus non-target-directed CoM (*ΔCoM)*. Positive *ΔCoM* values indicate that target-directed excursions were more likely to be followed by reversals towards the sound reference than non-target-directed excursions. We used linear mixed-effects models to examine main effects, with participant as a random intercept and audiovisual disparity as the main predictor. Candidate models included linear and quadratic effects of audiovisual disparity, with participant-specific motor noise considered as an additional covariate. Model fitting and post hoc contrasts followed the general statistical approach described above.

## Data and code availability

The data and materials necessary to replicate the current findings are available on the Radboud Data Repository (RDR); upon publication, a persistent identifier (DOI) will be made publicly available.

## Acknowledgement

This research was funded by the European Research Council (ERC Advanced Grant MakingSense 101096659).

---

**Figure S1.**
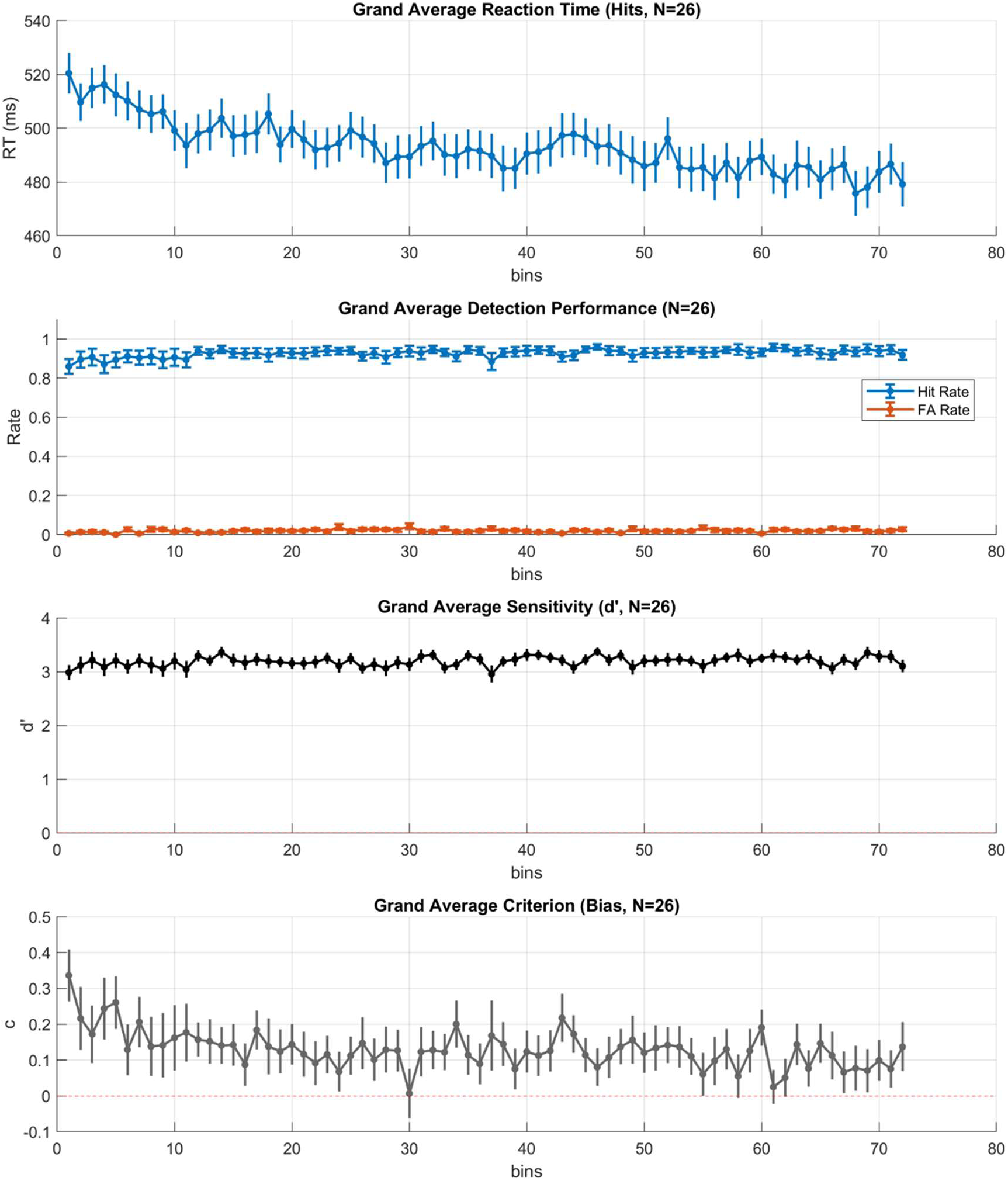
Phase A behavioural metrics. (A) Reaction time for hit trials across all Phase A bins in the retained behavioural sample (n = 26). Each bin corresponds to one short Phase A block of 31 trials. Dots indicate the across-participant mean for each bin, and error bars show ±SEM across participants. (B) Detection performance across the same bins. Hit and false-alarm rates were computed within each bin. Dots show across-participant means, and error bars show ±SEM across participants. (C) Signal-detection sensitivity (d′) across bins. d′ was calculated with a log-linear correction to avoid infinite z-scores. (D) Response criterion (c) across bins. Criterion was calculated with the same log-linear correction as d’; positive values indicate a more conservative response criterion. This figure provides a descriptive performance check rather than a separate group-level inferential test, confirming high hit rates, low false-alarm rates, stable positive sensitivity, and modest criterion fluctuations during the instrumental learning phase.

**Figure S2.**
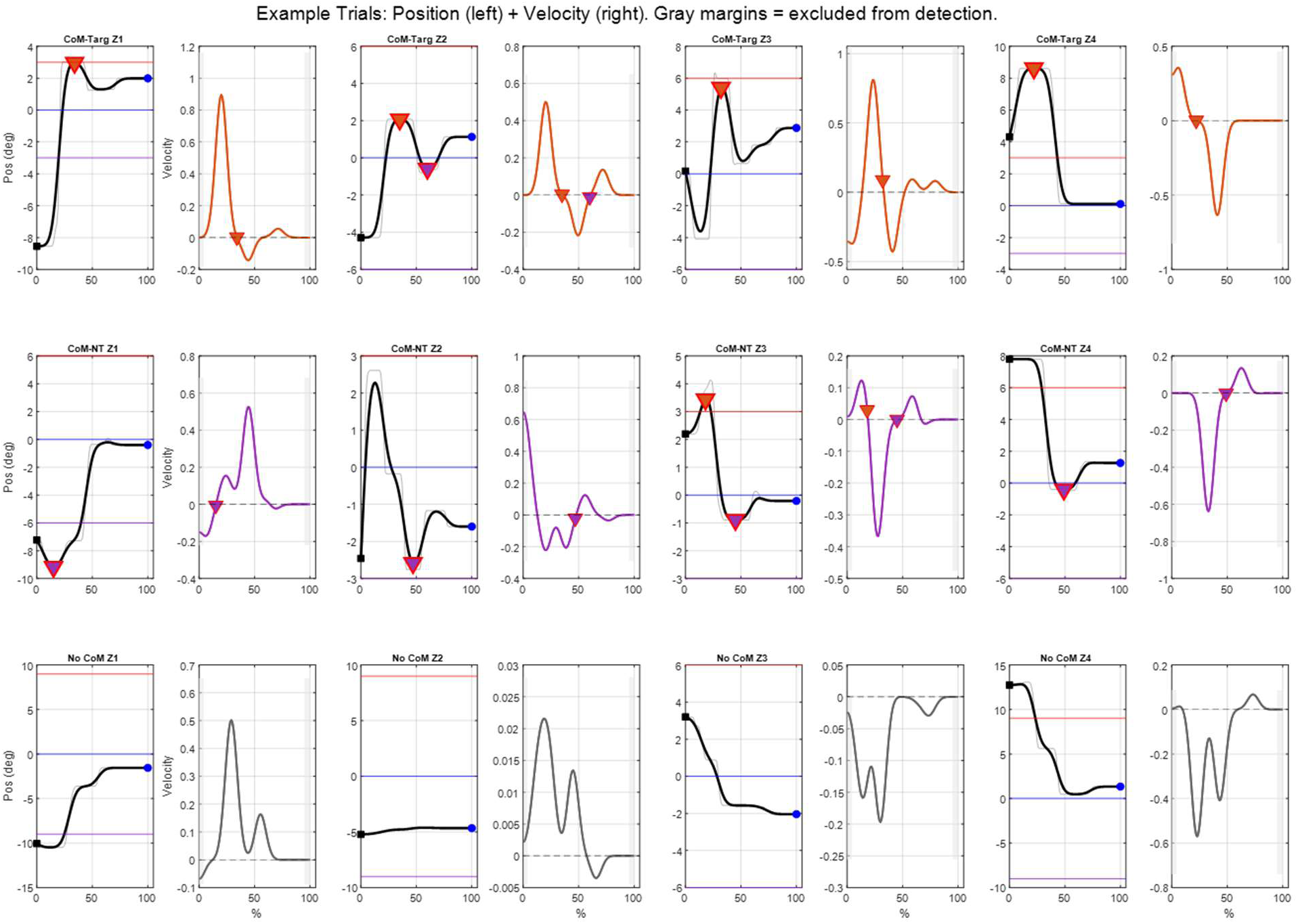
Velocity-based classification of CoM reversals. (A) Example target-directed CoM trials. Traces are shown after aligning the sound to 0° and flipping the spatial axis so that the target is on the positive side and the non-target is on the negative side. Columns show the four cursor starting zones relative to the non-target (NT), sound (S), and target (T): cursor-NT-S-T, NT-cursor-S-T, NT-S-cursor-T, and NT-S-T-cursor. Each zone is represented by a position panel on the left and a velocity panel on the right. In position panels, grey traces show the time-normalised raw cursor trajectory, black traces show the smoothed trajectory used for detection, the blue horizontal line marks the sound, the red line marks the target, and the purple line marks the non-target. Black squares mark cursor start positions, blue circles mark final cursor positions, and red inverted triangles mark detected reversal points. A target-directed CoM event was classified when a qualifying positive-velocity phase towards the target was followed by a qualifying negative-velocity phase back towards the sound. (B) Example non-target-directed CoM trials, using the same position and velocity display as in (A). These examples illustrate the mirrored version as target-directed conditions: a qualifying negative-velocity phase toward the non-target, followed by a qualifying positive-velocity reversal back toward the sound. (C) Example no-CoM trials, showing trajectories that did not satisfy the velocity, phase-duration, excursion, reversal-magnitude, and beyond-sound criteria required for CoM classification. In velocity panels, coloured traces show the velocity of the smoothed trajectory; dashed horizontal lines mark zero velocity, and grey shaded regions mark the start and end margins excluded from phase detection. This figure illustrates the classification procedure.

